# Comparative RNA-seq analysis of resistant and susceptible banana genotypes reveals molecular mechanisms in response to *Banana bunchy top virus* (BBTV)

**DOI:** 10.1101/2022.09.08.507103

**Authors:** Darlon V. Lantican, Jen Daine L. Nocum, Anand Noel C. Manohar, Jay-Vee S. Mendoza, Roanne R. Gardoce, Grace C. Lachica, Lavernee S. Gueco, Fe M. Dela Cueva

## Abstract

Banana is a major fruit crop in the Philippines and remains to be a large contributor to the country’s dollar reserve. Among the main hindrances in global banana production, diseases such as Banana bunchy top disease (BBTD) caused by BBTV can bring catastrophic loss to any banana plantation. To elucidate the resistance mechanism and understand the interplay of host factors in the presence of the invading pathogen, we implemented RNA-seq-based comparative transcriptomics analyses of mock- and BBTV-inoculated resistant (wild *M. balbisiana*) and susceptible (*M. acuminata* ‘Lakatan’) banana genotypes. Similar patterns of expression for 119 differentially expressed genes (DEGs) were observed on both genotypes, representing the typical defense response of banana to BBTV. A set of 173 DEGs specific to the susceptible ‘Lakatan’ banana cultivar revealed potential host factors and susceptibility mechanisms involved in successful BBTV infection. Further, differential transcriptomic analysis revealed 268 DEGs exclusive to the resistant wild *M. balbisiana*, unraveling insights into the complex resistance mechanisms involved in BBTV defense such as pathogen perception, phytohormone action, reactive oxygen species (ROS), hypersensitive response (HR), production of secondary metabolites and cell wall modification. The DEGs identified in this study will aid in the design of foreground markers for the precise integration of resistance genes during marker-assisted breeding programs. Furthermore, the application of these results will also enable the foreseen deployment of genome-edited banana cultivars targeting the resistance and host factor genes towards a future-proof banana industry.

## INTRODUCTION

Banana is a major fruit crop in the Philippines and remains to be a large contributor to the country’s dollar reserve, with an estimated total production of 2.40 million metric tons during the last quarter of 2020^1^. ‘Cavendish’, ‘Saba’ and ‘Lakatan’ varieties had the highest production of 50.4%, 28.5% and 10.7%, respectively. Among the 57 banana cultivars in the Philippines, the most planted varieties are ‘Saba’, ‘Latundan’, ‘Lakatan’, ‘Bangulan’ and ‘Cavendish’. The ‘Cavendish’ variety contributes to about 50% of the country’s total export, being highly preferred by the global market^2^. Nevertheless, the Philippines is still ranked 3^rd^ next to India and China in terms of 2010-2015 average global production but ranked 2^nd^ (next to Ecuador) in terms of 2013 agricultural export value^3^.However, the maximum potential in banana production is hindered by biotic and abiotic factors, such as the spread of destructive plant diseases like the banana bunchy top disease (BBTD).

BBTD, caused by *banana bunchy top virus (*BBTV), is transmitted by an aphid vector (*Pentalonia nigronervosa* Coquerel) and results to chlorotic and rosette leaves, stunted growth, and an inability to produce fruits^4,5^. BBTV transmission is circulative, non-propagative and persistent - the virus is non-replicative but remains in the aphid and transmits the virus through feeding into the phloem of one host to another. The infection also occurs in other banana planting materials, hence the faster spread of the disease^4^. One effective way to control BBTD is early detection of the vector and/or host infection and immediate replacement of infected plantlets with BBTV-free planting materials. Still, deployment of BBTV-resistant banana varieties in conjunction with a high-throughput, rapid and sensitive BBTV detection system is still the most sustainable approach to attaining optimum banana production. BBTV-resistant ‘Mapilak’ banana cultivar was previously developed through gamma irradiation and *in vitro* induced mutations of *Musa acuminata* ‘Lakatan’^6^. However, the resistance reaction of this cultivar only accounted for 20% of the disease incidence reduction^7^. Hence, identification of new sources of resistance alleles to be introgressed in the existing elite varieties is still an area of further research. Fortunately, the Philippines has local access to natural genetic resources such as wild *Musa balbisiana* accessions with complete resistance against the pathogen^8^.

Comparative RNA-seq analysis has aided in unraveling the mechanisms of disease resistance and identification of host-factor genes in several crops. The response mechanisms of banana during infection of tropical race 1 and 4 of *Fusarium oxysporium*^9^ and *Xanthomonas campestris* pv. *musacearum*^10^ were also effectively elucidated based on this approach. To date, there are still no publications or reports explaining the molecular mechanisms of banana-BBTV interaction with varying resistance and susceptibility responses.

With the availability of validated BBTV-resistant wild *M. balbisiana* genotype from the Philippines, identification of host-factors and resistance genes through transcriptomics studies (ie. RNA-seq) is now possible. Here, we provide insights into the molecular basis of disease resistance and host susceptibility between the BBTV-resistant genotype wild *M. balbisiana* and BBTV-susceptible genotype ‘Lakatan’.

## MATERIALS AND METHODS

### Preparation of biological samples and aphid-assisted inoculation

BBTV-resistant wild *M. balbisiana* (13-155) and BBTV-susceptible *M. acuminata* ‘Lakatan’ from the collections of the National Plant Genetic Resource Laboratory (NPGRL), Institute of Plant Breeding, University of the Philippines Los Baños (IPB-UPLB) were used for the transcriptomic analysis. For the BBTV-challenged treatment, three biological replicates each of wild *M. balbisiana* and *M. acuminata* ‘Lakatan’ were inoculated with BBTV by transferring 50 individual viruliferous aphids (*P. nigronervosa*) to each seedlings. On the other hand, the mock-inoculated treatment was prepared by feeding 50 individual BBTV-negative non-viruliferous aphids to each of the three (3) wild *M. balbisiana* seedlings and three (3) *M. acuminata* ‘Lakatan’ seedlings. RNA samples were extracted from young leaf tissues of banana seedlings 72 hours after inoculation (hpi).

### RNA Extraction of BBTV-resistant and BBTV-susceptible banana accessions

Total RNA from young leaf tissues of three biological replicates of BBTV-susceptible and BBTV-resistant *M. balbisiana* and *M. acuminata* ‘Lakatan’ (BBTV-inoculated and mock-inoculated) were extracted using the modified RNeasy Plant Mini Kit Protocol (QIAGEN, GmbH, Hilden, Germany). 200 mg of plant tissues from each banana samples were homogenized in liquid nitrogen in double sterilized mortar and pestle.

RNA concentration and quality were determined by gel electrophoresis in 1% UltraPure™ agarose (Invitrogen Corp., Carlsbad, California, USA) in 1X TBE running buffer at 100 V for 30 min. The gel was stained with 0.1 uL Gel Red (Biotium, CA, USA) visualized under UV illumination at 300 nm using the Enduro GDS Touch Imaging System (Labnet International, Inc, Edison, New Jersey, USA). RNA was quantified using Qubit™ RNA HS Assay Kit (Life Technologies, Thermo Fisher Scientific Inc.), Qubit™ 3.0 Fluorometer (Life Technologies, Thermo Fisher Scientific Inc.) following the manufacturer’s instruction.

### Outsourced next-generation sequencing of extracted RNA

High-quality RNA samples from wild *M. balbisiana* and *M. acuminata* ‘Lakatan’ were sent to the Philippine Genome Center – DNA Sequencing Core Facility (PGC-DSCF) for outsourced next-generation sequencing. Quality control check was done through RNA quantitation, 260/280 and 260/230 absorbance measurements, and gel separation using Microchip Electrophoresis System (MCE™-202 MultiNA; Shimadzu Biotech, Kyoto, Japan). Sequencing libraries were constructed using TruSeq Stranded Total RNA Library Prep Plant (Illumina), followed by sequencing on the Illumina NextSeq 500/550 sequencer.

### Bioinformatics analysis

The paired-end reads were mapped to the *M. acuminata* version 2 genome^11^ using STAR^12^ and then quantified using RSEM^13^, both at default settings. A matrix of the read count data generated from RSEM quantification was prepared and imported to R package DESeq2^14^ using tximport^15^.

A single-factor design was implemented to consider the factor introduced by the differences in the inoculation state (BBTV-inoculated vs. mock-inoculated) of the samples. After normalization using the relative log normalization (RLE) method, the DEGs between the two genotypes were identified. The Wald test from the general linear model (GLM) fitting was used to determine *p*-values. False discovery rate (FDR) values were derived using the Benjamini-Hochberg *p*-value adjustment algorithm. Genes with FDR values < 0.1 were considered differentially expressed between BBTV-inoculated and mock-inoculated ‘Lakatan’ and wild *M. balbisiana*. The gene set enrichment analysis (GSEA) with the GO data from the banana annotation files, which employs the Wallenius method to calculate the unbiased category enrichment scores, was performed using g:Profiler^16^.

## RESULTS

### Transcriptomes of resistant and susceptible banana genotypes in response to BBTV infection

Two banana genotypes, wild *M. balbisiana* (BBTV-resistant) and *M. acuminata* ‘Lakatan’ (BBTV-susceptible), were mock- and BBTV-inoculated by *P. nigronervosa* and the RNA samples from young leaf tissues were isolated at 72 hpi. In total, 12 RNA samples were sent for outsourced next-generation sequencing using Illumina NextSeq 500/550 to represent the total transcriptomic profiles of BBTV-free and BBTV-inoculated resistant and susceptible banana genotypes. Approximately 40-67 million raw reads (75-bp length per read) were generated from each sequencing library. All the raw reads generated from this study were uploaded to NCBI under BioProject Accession number PRJNA746416.

The *M. acuminata* version 2 genome assembly^11^ from the Banana Genome Hub (https://banana-genome-hub.southgreen.fr) was used as reference in the genome-guided mapping of the RNA-seq reads for subsequent differential gene expression transcriptomic analysis. Around 18 – 30 million paired-end reads (150 bp) of the RNA-seq data were observed to uniquely map to the reference genome. On average, 93.46% (mock-inoculated ‘Lakatan’), 92.74% (BBTV-inoculated Lakatan), 92.61% (mock-inoculated wild *M. balbisiana*), and 93.59% (BBTV-inoculated wild *M. balbisiana*) were mapped to the genome (Supplementary Table S1).

### Changes in expression patterns between wild M. balbisiana (resistant) and ‘Lakatan’ (susceptible) in response to BBTV

The RSEM-normalized expected count data from BBTV-challenged wild *M. balbisiana* (24,670 genes with non-zero read count) and ‘Lakatan’ (27,285) were statistically compared using the DESeq2 R package^14^. The DESeq2 pipeline was able to identify 292 and 387 differentially expressed genes upon BBTV-infection in ‘Lakatan’ and wild *M. balbisiana*, respectively (Supplementary Table S2). Among these significantly expressed genes, 134 genes were up-regulated while 158 genes were down-regulated in ‘Lakatan’, the susceptible banana cultivar. Meanwhile, the BBTV-resistant wild *M. balbisiana* was observed to up-regulate 123 genes and down-regulate 264 genes (Volcano plot; Figure 1). Moreover, comparison of the differentially expressed gene (DEG) profiles between the susceptible and resistant banana accessions revealed 268 DEGs detected only on the resistant wild banana (Venn Diagram; Figure 1). Results further showed that the susceptible ‘Lakatan’ banana variety differentially expressed 173 genes, exclusively. This set of genes unique to ‘Lakatan’ may include host factors necessary for a successful BBTV infection in banana. In addition to these DEGs, similar profiles for 119 DEGs are observed on both genotypes, suggesting their possible roles in the recognition and basal defense response mechanism of banana upon BBTV introduction.

**Figure 1.**
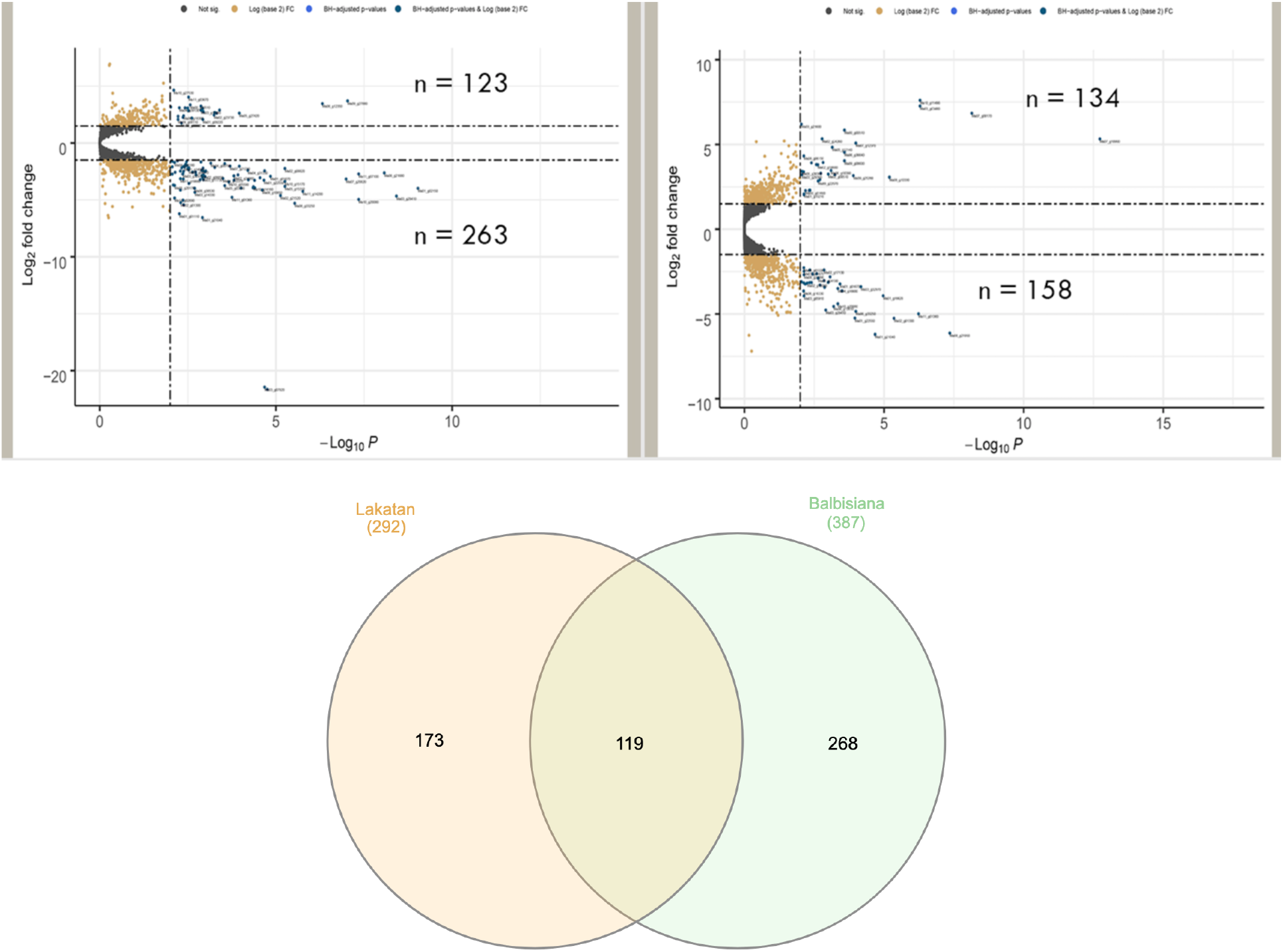
Volcano plots and Venn diagram of DEGs. Volcano plots of differentially expressed genes (DEGs) in wild *Musa balbisiana* (**A**) *Musa acuminata* ‘Lakatan’ (**B**) three (3) days after BBTV inoculation. The y-axis shows the fold change difference in the expression of genes, and the x-axis indicates the BH-adjusted p-values for the changes in the expression. An absolute value of log2 fold change >1 and the *p*-value < 0.1 was set to declare differentially expressed genes (DEGs). (**C**) Venn diagram showing the overlap of DEGs between wild *M. balbisiana* and *M. acuminata* ‘Lakatan’.

### Shared set of DEGs in resistant and susceptible banana cultivars involved in defense response

Among the identified DEGs, 119 were present in both cultivars in response to BBTV infection (Supplementary Table S2). These includes DEGs involved in pathogen response such as cysteine-rich receptor-like kinase 6 (CRK6) (Ma06_g23640), protein kinase phototropin 1A (Ma04_g37140), calcium-dependent protein kinase (CDPK) (Ma06_g31170), mitogen-activated protein kinase (MAPK) (Ma09_g06560), and mitogen-activated protein kinase kinase kinase (MAPKKK)(Ma08_g21950). The gene expression patterns of these genes were the same between wild *M. balbisiana and M. acuminata*, in which CRK6, CDPK, and phototropin 1A were up-regulated while MAPK and MAPKKK were down-regulated. CRKs generally plays a role against pathogen attack and programmed cell death^17–20^; specifically, CRK6 is involved in signaling in response to extracellular reactive oxygen species (ROS)^21^. Phototropin 1A is a member of AG{Bibliography}C-VIII kinases which function as photoreceptors, regulates auxin transport and contribute to pathogen response^22–24^. MAPK, MAPKKK and CDPK regulate plant immune signaling enabling activation of defense genes expression^25,26^. On the other hand, heat shock proteins (HSPs) chaperone HSP70 (Ma06_g36160) and co-chaperone HSP40/DnaJ (Ma04_g37990) were observed to be both downregulated in the two banana genotypes in response to BBTV. These HSPs generally act as chaperones in pathogenesis, particularly movement, demonstrating their possible role in plant defense or immunity^27^.

Thirteen (13) differentially expressed transcription factors (TFs) were all observed to be downregulated in both cultivars upon BBTV introduction which comprise of two B-box (BBX) (Ma10_g08680, Ma04_g37190), three Dof (Ma06_g33890, Ma09_g04960, Ma11_g14200), six MYB (Ma06_g16330, Ma10_g14160, Ma01_g01670, Ma02_g01300, Ma04_g16680, Ma11_g01360), a C2H2 (Ma03_g14330) and a bHLH (Ma08_g10850) TFs.

The importance of these transcription factors in defense response of the plant host against pathogen invasion has been previously established in other crops^28–34^.

### Differential expression of genes specific to susceptible cultivar

*M. acuminata*-specific DEGs were further investigated to shed light on the response of a susceptible genotype during BBTV invasion as well as identify potential host factors needed by the pathogen for a successful disease progression. During BBTD development, DEGs involved in the biogenesis of large ribosomal subunit 60S (Ma07_g21640) and small ribosomal subunit 40S (Ma01_g15210, Ma07_g07360) were up-regulated in the susceptible cultivar. Expression of translation elongation factor eEF1A (Ma_ 10_g13420) was likewise increased (~2x fold) in response to the invading virus. Being a critical step for successful viral replication, subversion of the host’s protein synthesis machinery and increasing the expression of the components, such as the eEF1A, are needed by the viral pathogen^35^. Genes involved in RNA biosynthesis and processing such as RNA polymerase sigma factor sigE (Ma07_g26010), group-II intron splicing RNA helicase RH3 (Ma11_g12630), and phosphorolytic exoribonuclease PNP (Ma10_g20770) were also up-regulated in the susceptible variety, signifying their potential involvement in BBTV replication.

Interestingly, seven HSPs involved in protein homeostasis were up-regulated in the *M. acuminata* ‘Lakatan’ comprising of mitochondrial heat shock 70 kDa protein (mtHsc70; Ma04_g26470), HSP70 (Ma02_g18000), HSP40 (Ma04_g29330), HSP90 (Ma03_g29390), Hsp60-co-chaperone Hsp20 (Ma05_g04870), and two homologs of HSP60 (Ma09_g23670, Ma11_g20430). The heat shock proteins available in the host are needed for the virus’ mechanisms for protein processing and have an impact on viral proliferation and counteracting the host’s defense responses^36–38^. Moreover, gene expression of two homologs of vesicle-associated membrane protein (VAMP)-associated proteins (VAPs) (Ma07_g07110; Ma11_g08690) and an NAD-dependent glyceraldehyde 3-phosphate dehydrogenase (Ma06_g01470) were also upregulated. These genes from the host were previously reported to be implicated in the viral cell-to-cell movement and replication^39–41^. Hence, the observed upregulation of these sets of DEGs plays a direct role in the development of BBTD in the susceptible banana cultivar.

Genes involved in phytohormone action, specifically, auxin signaling and jasmonic acid biosynthesis, were also differentially expressed in *M. acuminata* ‘Lakatan’. Regulation of these phytohormones during plant-pathogen interaction is critical for plant susceptibility or defense response^42–44^. The IAA/AUX repressor (Ma09_g10330) regulating the phytohormone auxin was down-regulated by ~1.5-fold while the allene oxidase synthase driving the biosynthesis of jasmonic acid was up-regulated by ~3-fold in banana in response to BBTV infection. The pathogenesis-related protein 1 (PR1, Ma08_g34160) related to the salicylic-acid mediated resistance was also observed to be upregulated in the susceptible cultivar by ~2.3-fold.

Notably, two genes integral to the host SCF (SKP1-CUL1-F-box protein) ubiquitin ligase complex, substate adaptor BT (Ma11_g13780) and substrate adaptor ADO (Ma07_g10990), were both over-expressed in BBTV-infected *M. acuminata* with ~2.3-fold change and ~5.3-fold-change, respectively. The SCF ubiquitin ligase complex, which plays a key role in host defense response, has been reported to be hijacked by invading viruses for their own advantage^45^.

### Differential expression of defense-associated genes in resistant cultivar

Two-hundred sixty-eight (268) DEGs were found to be specific to the BBTV-resistant wild *M. balbisiana* which will provide insights into the molecular mechanisms of resistance to overcome BBTD disease progression. Among the set of DEGs specific to the resistant cultivar, nine protein kinases were identified, which may play an essential role in pathogen recognition and signaling for a cascade of plant defense mechanisms^46^. Interestingly, the wall-associated kinase (WAK/WAKL; Ma05_g19150), involved in the maintenance of cell wall integrity under stress^47^, was over-expressed by ~3-fold in the resistant cultivar in response to pathogen attack. Leucine-Rich Repeat (LRR) protein kinases LRR-VI (Ma07_g26720; downregulated gene expression by ~2.5-fold) and LRR-VIII-2 (Ma08_g06820; up-regulated gene expression by ~1.2-fold), which can contribute to activating defense/basal response against BBTV^48–50^, were differentially expressed in wild *M. balbisiana*. The gene expression of hypersensitive induced response protein 1 (Ma07_g25320) and hypersensitive-induced response protein-like protein 1 (Ma07_g25340) in resistant banana cultivar were also observed to respond to BBTV invasion with increased gene expression by ~1.4-fold and ~1.6-fold compared to the mock-inoculated control, respectively.

Plant immune system is also regulated by the action of phytohormones, which synergistically and/or antagonistically work in a complex network to respond to the invading pathogen^51,52^. The genes encoding nematode resistance protein HSPRO2 (Ma10_g05760) and nematode resistance protein-like HSPRO2 (Ma11_g22440), which play roles in hormone-mediated stress signaling^53^, were both downregulated in resistant banana cultivar. Differential gene expression in wild *M. balbisiana* was also observed in the perception modulator ABAR (Ma02_g08820; downregulated by ~2.2-fold) involved in abscisic acid signaling. Downregulation of gene expression in auxin transport genes such as auxin efflux carrier component 1a (Ma08_g23810) and auxin efflux carrier component 5 (Ma06_g01320) were also detected as a response of BBTV-resistant cultivar to the invading viral pathogen.

Changes in the reduction-oxidation (redox) levels signal defense response mechanisms in pathogen-challenged plant host cells^54^. In the BBTV-challenged resistant banana cultivar, the thioredoxin-domain containing protein (Ma08_g22250) associated with thiol-based redox regulation was downregulated by ~2.8-fold. Similarly, genes implicated in ascorbate-based redox regulation, such as cyt-b561 electron shuttle hemoprotein CYBASC (Ma01_g02150) and apoplastic ascorbate oxidase AAO (Ma07_g08950), were shown to be downregulated in wild *M. balbisiana*.

The resistant response of wild *M. balbisiana* to BBTV was also accompanied by differential expression of genes for cell wall modifying enzymes. These genes include xylan alpha-1,3-arabinosyltransferase (Ma02_g24680; down-regulated by ~3.06) for xylan biosynthesis; glucuronoxylan 4-O-methyltransferase (Ma08_g29650; up-regulated by ~3.61) for xylan modification and degradation, hydroxycinnamate glucosyltransferase (Ma01_g00430; down-regulated by ~2.83) for cell wall hydroxycinnamic acids, and two homologs of caffeic acid O-methyltransferase (Ma01_g01680 downregulated by ~1.40; Ma01_g01680 up-regulated by ~2.61) for monolignol biosynthesis.

Genes involved in secondary metabolite production, such as terpenoids and phenolics, were also shown to be differentially expressed in the resistant cultivar. Specifically, three genes involved in terpenoid biosynthesis [4-hydroxy-3-methylbut-2-enyl diphosphate reductase (Ma08_g04770), phytoene synthase (Ma09_g09640), carotenoid beta-ring hydroxylase BCH (Ma11_g19880)] were downregulated while two gene homologs of (-)-alpha-terpineol synthase (TPS; Ma04_g07700, Ma04_g08150) were up-regulated. On the other hand, the type-I flavone synthase (FS; Ma02_g12040) which catalyzes the conversion of flavanones to flavones was downregulated in the BBTV-resistant banana cultivar upon BBTV infection by ~3.4-fold. Genes involved in polyamine biosynthesis were differentially expressed in the resistant cultivar in response to the viral invasion. Specifically, gene expression of arginine decarboxylase (Ma09_g21350) associated with putrescine biosynthesis and S-adenosyl methionine decarboxylase (Ma05_g26570) which participates in spermidine biosynthesis were both downregulated by ~1.42 and ~1.39, respectively.

### Gene ontology (GO) enrichment of differentially expressed genes

To investigate the functionality of the genes activated in response to BBTV infection in wild *M. balbisiana* as compared to that in the susceptible ‘Lakatan’ cultivar, the direct counts of GO terms associated with the differentially expressed genes were assessed (Figure 2). The top overrepresented Biological Process (BP) GO terms based on the differentially expressed genes (DEGs) upon BBTV infection in both the banana cultivars are involved ‘cellular process’, ‘metabolic process’, ‘organic substance metabolic process’, ‘cellular metabolic process’ and ‘primary metabolic process’. For Molecular Function (MF), GO terms associated with ‘binding’, ‘catalytic activity’, ‘organic cyclic compound binding’ and ‘heterocyclic compound binding’ were observed to have the most enriched MF GOs. Lastly, analysis of the Cellular Component (CC) GO terms in DEGs in both *M. acuminata* ‘Lakatan’ and wild *M. balbisiana* revealed high representation for ‘cellular anatomical entity’, ‘intracellular anatomical organelle’, ‘intracellular organelle’, ‘organelle’ and ‘cytoplasm’. Generally, the direct count of GO terms (BP, MF, and CC) was always higher in wild *M. balbisiana* than in *M. acumanata* “Lakatan”, except for the CC GO term ‘nucleus’ in which *M. acuminata* “Lakatan” has a higher number of associated DEGs (19 DEGs) than wild *M. balbisiana* (14 DEGs).

**Figure 2.**
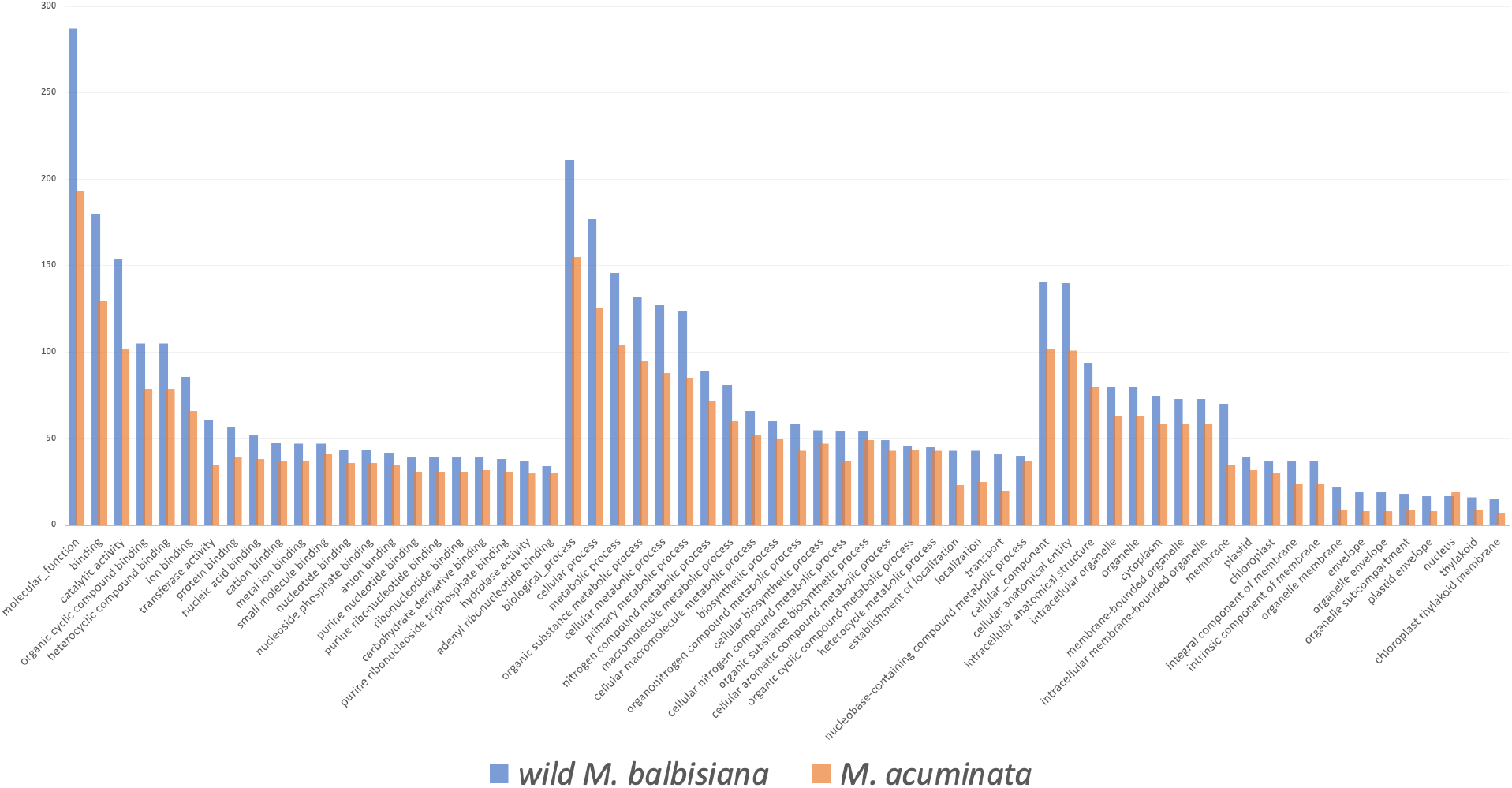
Direct counts of the over-represented GOs in wild M. balbsiana and ‘Lakatan’ in response to BBTV inoculation.

Gene Ontology (GO) and Kyoto Encyclopedia of Genes and Genomes (KEGG) enrichment analyses (Table 1) were carried out to further identify which biological processes the differentially expressed gene sets in both genotypes were enriched in and what detailed metabolic pathways they engaged in using g:Profiler^16^. Four significantly enriched MF GO terms were found in ‘Lakatan’ [GO:0004645; GO:0008184; GO:0102250; GO:0102499], compared to seven in wild *M. balbisiana* [GO:0016157; GO:0008519; GO:0035251; GO:0008194; GO:0102499; GO:0102250; GO:0008184]. Significant enrichment of MFs was identified for the GO terms GO:0008184, GO:0102250, and GO:0102499 in both genotypes. The 1,4-alpha-oligoglucan phosphorylase activity (GO:0004645), in contrast, was only found in ‘Lakatan’, while the transporter activities and sucrose synthase activity (GO:0016157, GO:0008519, GO:0035251, and GO:0008194) were only present in the BBTV-resistant wild *M. balbisiana*.

**Table 1.** Significant GO terms (MF, BP and CC) and KEGG pathways based on the differentially expressed genes (DEGs) in BBTV-susceptible and -resistant genotypes.

| Banana Genotype | Source | GO Name | GO ID | adjusted p value | negative log10 of adjusted p value | Term size | Query size | Intersection size |
| --- | --- | --- | --- | --- | --- | --- | --- | --- |
| Lakatan | GO:MF | 1,4-alpha-oligoglucan phosphorylase activity | GO:0004645 | 0.0153 | 1.8160 | 8 | 198 | 3 |
| Lakatan | GO:MF | glycogen phosphorylase activity | GO:0008184 | 0.0289 | 1.5388 | 2 | 198 | 2 |
| Lakatan | GO:MF | linear malto-oligosaccharide phosphorylase activity | GO:0102250 | 0.0289 | 1.5388 | 2 | 198 | 2 |
| Lakatan | GO:MF | SHG alpha-glucan phosphorylase activity | GO:0102499 | 0.0289 | 1.5388 | 2 | 198 | 2 |
| Lakatan | GO:BP | response to light stimulus | GO:0009416 | 0.0008 | 3.1026 | 213 | 152 | 12 |
| Lakatan | GO:BP | response to radiation | GO:0009314 | 0.0013 | 2.8743 | 224 | 152 | 12 |
| Lakatan | GO:BP | response to abiotic stimulus | GO:0009628 | 0.0049 | 2.3134 | 443 | 152 | 16 |
| Lakatan | GO:BP | circadian rhythm | GO:0007623 | 0.0077 | 2.1141 | 32 | 152 | 5 |
| Lakatan | GO:BP | rhythmic process | GO:0048511 | 0.0139 | 1.8558 | 36 | 152 | 5 |
| Lakatan | GO:CC | plastid | GO:0009536 | 0.0000 | 5.5471 | 883 | 96 | 26 |
| Lakatan | GO:CC | chloroplast | GO:0009507 | 0.0000 | 4.6704 | 843 | 96 | 24 |
| Lakatan | KEGG | Biosynthesis of cofactors | KEGG:01240 | 0.0380 | 1.4207 | 111 | 26 | 6 |
| wild *M. balbisiana* | GO:MF | sucrose synthase activity | GO:0016157 | 0.0032 | 2.4984 | 11 | 292 | 4 |
| wild *M. balbisiana* | GO:MF | ammonium transmembrane transporter activity | GO:0008519 | 0.0047 | 2.3273 | 12 | 292 | 4 |
| wild *M. balbisiana* | GO:MF | UDP-glucosyltransferase activity | GO:0035251 | 0.0284 | 1.5469 | 76 | 292 | 7 |
| wild *M. balbisiana* | GO:MF | UDP-glycosyltransferase activity | GO:0008194 | 0.0418 | 1.3783 | 270 | 292 | 13 |
| wild *M. balbisiana* | GO:MF | SHG alpha-glucan phosphorylase activity | GO:0102499 | 0.0499 | 1.3016 | 2 | 292 | 2 |
| wild *M. balbisiana* | GO:MF | linear malto-oligosaccharide phosphorylase activity | GO:0102250 | 0.0499 | 1.3016 | 2 | 292 | 2 |
| wild *M. balbisiana* | GO:MF | glycogen phosphorylase activity | GO:0008184 | 0.0499 | 1.3016 | 2 | 292 | 2 |
| wild *M. balbisiana* | GO:BP | cellular carbohydrate metabolic process | GO:0044262 | 0.0013 | 2.8957 | 319 | 210 | 17 |
| wild *M. balbisiana* | GO:BP | ammonium transmembrane transport | GO:0072488 | 0.0062 | 2.2108 | 11 | 210 | 4 |
| wild *M. balbisiana* | GO:BP | response to abiotic stimulus | GO:0009628 | 0.0075 | 2.1251 | 443 | 210 | 19 |
| wild *M. balbisiana* | GO:BP | sucrose metabolic process | GO:0005985 | 0.0172 | 1.7636 | 27 | 210 | 5 |
| wild *M. balbisiana* | GO:BP | carbohydrate metabolic process | GO:0005975 | 0.0206 | 1.6856 | 1032 | 210 | 31 |
| wild *M. balbisiana* | GO:BP | response to light stimulus | GO:0009416 | 0.0239 | 1.6222 | 213 | 210 | 12 |
| wild *M. balbisiana* | GO:BP | response to radiation | GO:0009314 | 0.0388 | 1.4107 | 224 | 210 | 12 |
| wild *M. balbisiana* | GO:CC | plastid thylakoid membrane | GO:0055035 | 0.0000 | 4.6784 | 143 | 134 | 12 |
| wild *M. balbisiana* | GO:CC | chloroplast thylakoid membrane | GO:0009535 | 0.0000 | 4.6784 | 143 | 134 | 12 |
| wild *M. balbisiana* | GO:CC | plastid | GO:0009536 | 0.0000 | 4.5796 | 883 | 134 | 30 |
| wild *M. balbisiana* | GO:CC | plastid thylakoid | GO:0031976 | 0.0000 | 4.4657 | 179 | 134 | 13 |
| wild *M. balbisiana* | GO:CC | chloroplast thylakoid | GO:0009534 | 0.0000 | 4.4657 | 179 | 134 | 13 |
| wild *M. balbisiana* | GO:CC | chloroplast | GO:0009507 | 0.0001 | 3.9377 | 843 | 134 | 28 |
| wild *M. balbisiana* | GO:CC | thylakoid membrane | GO:0042651 | 0.0002 | 3.7581 | 174 | 134 | 12 |
| wild *M. balbisiana* | GO:CC | thylakoid | GO:0009579 | 0.0003 | 3.5437 | 251 | 134 | 14 |
| wild *M. balbisiana* | GO:CC | plastid membrane | GO:0042170 | 0.0005 | 3.2994 | 227 | 134 | 13 |
| wild *M. balbisiana* | GO:CC | photosynthetic membrane | GO:0034357 | 0.0006 | 3.2406 | 195 | 134 | 12 |
| wild *M. balbisiana* | GO:CC | plastid envelope | GO:0009526 | 0.0147 | 1.8317 | 313 | 134 | 13 |
| wild *M. balbisiana* | KEGG | Metabolic pathways | KEGG:01100 | 0.0118 | 1.9269 | 992 | 40 | 29 |
| wild *M. balbisiana* | KEGG | Starch and sucrose metabolism | KEGG:00500 | 0.0125 | 1.9037 | 57 | 40 | 6 |
| wild *M. balbisiana* | KEGG | Glyoxylate and dicarboxylate metabolism | KEGG:00630 | 0.0371 | 1.4307 | 47 | 40 | 5 |

In response to BBTV invasion, the wild *M. balbisiana* genotype and the ‘Lakatan’ genotype significantly enriched five and seven BP GO terms, respectively. The enriched biological processes that both genotypes shared were response to light stimulus (GO:0009416), response to radiation stimulus (GO:0009314), and response to abiotic stimulus (GO:0009628). As such, cellular carbohydrate metabolic process (GO:0044262), ammonium transmembrane transport (GO:0072488), sucrose metabolic process (GO:0005985), and carbohydrate metabolic process (GO:0005975) were exclusive to wild *M. balbisiana*, whereas circadian rhythm (GO:0007623) and rhythmic process (GO:0048511) were only significantly enriched in ‘Lakatan’.

Plastid (GO:0009536) and chloroplast (GO:0009507) CC GOs were enriched in ‘Lakatan’. These CC GOs were also observed in wild *M. balbisiana* in addition to nine CC GO terms (GO:0055035; GO:0009535; GO:0009536; GO:0031976; GO:0009534; GO:0009507; GO:0042651; GO:0009579; GO:0042170; GO:0034357; GO:0009526) associated with thylakoid, thylakoid membrane, photosynthetic membrane, and plastid envelope in general. Based on KEGG enrichment, biosynthesis of cofactors (KEGG:01240) was the only identified significant pathway in the BBTV-susceptible ‘Lakatan’ genotype. On the other hand, three KEEG pathways (KEGG:01100 - metabolic pathway; KEGG:00500 - starch and sucrose metabolism; KEGG:00630 - glyoxylate and dicarboxylate metabolism) were significant in the BBTV-resistant wild *M. balbisiana* as part of the defense response against the pathogen.

## DISCUSSION

Almost all regions cultivating bananas are affected by the prevalence of banana bunchy top disease caused by BBTV. Eliminating infected plants and using virus-free tissue culture-derived planting materials can help to reduce the spread of the pathogen. Still, utilizing natural sources of resistance in breeding programs is still far the most effective and long-term strategy to combat its negative impact on the banana industry. Cultivated banana types, such as ‘Cavendish’ and ‘Lakatan’ varieties, are susceptible to BBTV^6,55^, unlike the wild type banana progenitor *M. balbisiana* which exhibits resistance to the disease^56^.

In recent years, the progress of next-generation sequencing and bioinformatics approaches, such as transcriptome-wide expression studies using RNA-seq data, has been heavily utilized in the study of plant-pathogen interactions. Such scientific undertakings can help in the development of molecular breeding strategies^57^ to mitigate the large-scale destructive effects of plant pathogens. Hence, we investigated the underlying molecular mechanisms of successful BBTD progression in a susceptible commercial ‘Lakatan’ banana variety in comparison with the resistance response of wild *M. balbisiana*. By using this approach, we were able to identify 387 DEGs in the BBTV-resistant banana genotype and only 292 in the BBTV-susceptible ‘Lakatan’ banana 72 hours after post-inoculation. A similar trend was also previously observed in the transcriptomic investigation of the response of *M. balbisiana* to *Xanthomonas campestris* pv. *musacearum* causing banana *Xanthomonas* wilt disease, in which the resistant *M. balbisiana* had higher number of identified DEGs as compared to the susceptible genotype, Pisang Awak^10^. In this work, differentially expressed gene sets were unraveled for the very first time, shedding insights on the underlying mechanisms controlling banana susceptibility and resistance to BBTV.

### Insights into typical defense response of banana against BBTV

Regardless of the banana genotype, 119 genes are commonly differentially expressed and have similar patterns of gene expression upon the exposure of the host plant to the invading pathogen. The defense reaction of the banana to BBTV, independent of whether a disease would arise or not, is represented by this shared gene set of both resistant and susceptible banana genotypes. To combat against invading pathogens, plants have many layers of defense^58^. When plants activate their defense mechanisms in response to a pathogen attack, plant-pathogen interactions occur^59^. Recognition of pathogen invasion is essential for triggering an effective plant defense response. To achieve this, plants possess pattern recognition receptors (PRRs) to recognize the evolutionary conserved pathogen’s molecular fingerprints known as pathogen-associated molecular patterns (PAMPs) or microbe-associated molecular patterns (MAMPs) resulting in pattern-triggered immunity (PTI)^60–63^. All known plant PRRs are categorized as either receptor-like kinases (RLKs) or receptor-like proteins (RLPs) with functional domains, both of which are present in the plasma membrane^64^. In Arabidopsis, cysteine-rich receptor like kinase 6 (*CRK6*) overexpression increased the PTI response and developed resistance to *Pseudomonas syringae* pv. tomato DC3000 by associating with a PRR, the FLAGELLIN SENSING2 (*FLS2*)^65^. CaCRK6 was also observed to heterodimerized with CaCRK5, suggesting its role in the innate immune response of pepper against *Ralstonia solanacearum* infection^66^. Hence, the up-regulation of *CRK6* (Ma06_g23640) in both banana genotypes may indicate its potential function in the recognition of BBTV to initiate basal defense response.

Following the recognition of the pathogen via RLKs or RLPs, plants activate downstream signaling networks regulated by Ca^2+^-dependent protein kinase (CDPK) and mitogen-activated protein kinase (MAPK) cascades^67^. To trigger secondary and late defensive responses, MAPK activations may need to exceed a threshold in both duration and amplitude and interact differentially or synergistically with Ca^2+^ signaling^67,68^. The activities and synthesis of various transcription factors (TFs), enzymes, hormones, peptides, and antimicrobial chemicals are regulated by MAPK and CDPK signaling networks in specific and overlapping manner that contribute to the defense response against various pathogens^67,69–71^. Thus, the down-regulation of two MAPKs (Ma09_g06560, Ma08_g21950) and the up-regulation of a CDPK (Ma06_g31170) gene expression in banana following the introduction of BBTV indicates a possible role in the regulation of processes essential to the defense response through kinase signaling pathways.

Light perception also contributes to the response of plants to pathogen attack by activating signaling pathways, regulating gene expression and controlling the cell death response^72,73^. To sense light, plants utilize a group of proteins collectively known as photoreceptors^74^. Interestingly, both resistant and susceptible banana genotypes significantly up-regulate the expression of the phototropin 1A (*PHOT1A*) gene, a photoreceptor for the perception of blue-light^75^. In Arabidopsis, HYPERSENSITIVE RESPONSE TO TCV (HRT)-mediated plant defense against the Turnip Crinkle Virus (TCV) directly depends on blue-light photoreceptors, PHOT1 and PHOT2^73^. This process is negatively regulated by CONSTITUTIVE PHOTOMORPHOGENIC 1 (COP1), a primary regulator of light signaling, by interacting and tagging HRT protein for degradation via the 26S proteosome. The interaction between COP1 and SUPPRESSOR OF PHYA-105 (SPA) proteins determines the E3 ligase activity of COP1 in plants^76^. In this study, the two banana genotypes used were observed to have downregulation of the COP1-SPA adaptor subcomplex COP1 (Ma06_g36160) and two homologs of regulator component SPA (Ma01_g07220.1; Ma03_g32970.1). Hence, the upregulation of *PHOT1A* gene and downregulation of the components of its potential key regulator, COP1-SPA E3 Ubiquitin Ligase complex, suggest the potential interaction among these genes to help in the defense of banana against BBTV.

The possible contribution of some HSPs to defend against the invading BBTV pathogen was also observed in both banana genotypes in which the expression of chaperone *HSP70* (Ma06_g36160) and co-chaperone *HSP40* were both observed to be downregulated in ‘Lakatan’ and wild *M. balbisiana*. In other plant species, upregulation of *HSP70* was observed as a molecular mechanism in response to pathogens such as *Phytophthera parasitica* in tomato^77^ and *Blumeria graminis* in barley^78^. This is also consistent with the observed overexpression of *HSP40* in soybean and tomato, demonstrating its role in disease resistance^79,80^.

### RNA-seq reveals putative susceptibility factors for BBTD progression in banana

Viruses heavily rely on the host’s machinery to promote the synthesis of vital components for viral replication and systemic cell-to-cell movement towards disease progression by hijacking various cellular processes^81,82^. For instance, viruses tamper with the host translation system by preventing the recruitment of cap-dependent ribosomes for the host’s mRNA translation (also known as “host shut-off”). Such viral strategy has an overall impact on the host’s anti-viral defense mechanism as well as promotes the synthesis of proteins for viral proliferation^35,83^. A similar strategy was also implemented by the invading BBTV pathogen during its course of infection in ‘Lakatan’ banana cultivar based on the results of our current transcriptomic investigation. Components of the banana protein biosynthesis system such as 40S ribosomal protein SA, rRNA biogenesis protein/40S assembly factor RRP5, glutamine- and glycine-tRNA ligases, elongation factor 1α, and 60S processome rRNA methyltransferase were all observed to be recruited by the invading pathogen to promote the biosynthesis of its own proteins for successful subversion of the host’s cells. Similar findings were obtained in a previous comparative transcriptomics study following cucumber mosaic virus (CMV) infection in the susceptible cucumber genotype, ‘Vanda’, where structural components of the ribosome and the translation biological process were substantially enriched and up-regulated^84^.

Viruses not only recruit plant translational components but also exploit and re-localize host genes involved in DNA replication and transcription to increase viral replication and spread^85,86^. Consequently, genes in banana genome implicated in the RNA biosynthesis and processing are somewhat being mobilized by BBTV for its advantage, as evidenced by the ‘Lakatan’-specific overexpression of DNA-dependent RNA polymerase I complex subunit NRPA2, RNA polymerase sigma factor sigE, group-II intron splicing RNA helicase RH3, and phosphorolytic exoribonuclease PNP. ssDNA viruses, such as BBTV, use the host’s DNA replication system to convert their ssDNA to intermediate dsDNA, which is subsequently transcribed into mRNA^87,88^. Moreover, the RNA processing of the host cell, such as splicing machinery, is needed by the virus for the maturation of its mRNA necessary for proliferation, survival, and adaptation within the host^88^. In relation to the increased activity in the host’s DNA replication and transcription system, the nucleotide metabolism in banana is being enhanced during BBTV infection supported by the up-regulation of the expression of genes involved in purine anabolism (aminoimidazole RN synthetase PUR5), pyrimidine phosphotransfers activity (UMP kinase PUMPKIN) and the deoxynucleotide salvage pathway (deoxyribonucleoside kinase TK2). Increased nucleotide metabolism is a metabolic change linked to a variety of viral infections^89,90^. Previous study demonstrates the interaction between host proteins involved in nucleotide biosynthesis and *Papaya ringspot virus* (PRSV) coat protein (CP) during infection in papaya^91^, signifying the potential role of these genes in disease progression in susceptible hosts.

Virulent pathogens must overcome the initial layer of resistance of the host for successful infection, leading to disease development. In a bid to subdue PTI, these pathogens introduce effector proteins into the plant cell. With this mechanism, pathogens can survive and complete their life cycle, developing effector-triggered susceptibility (ETS) of the host^63,92–94^. Plant virus effectors target various host cellular processes such as transcription factors regulating defense responses, protein degradation pathways, and re-localization of plant proteins^94^. Transcription factors classified as BBX-DBB, BBX-CO, DOF, MYB, REVEILLE, bZIP, NPL, ERF, AP2, NAC, and bHLH were differentially expressed in the susceptible cultivar during the BBTV infection process. Although no direct link has been established on how BBTV utilizes these host TFs via effectors, previous research shows that the association of betasatellites (βC1) with *Tomato yellow leaf curl China virus* (TYLCCNV) increased viral pathogenicity by binding to several transcription factors that regulate plant defense responses^95^. Recently, it was reported that a self-replicating alphasatellite associated with BBTV contributes to viral replication, transcription, siRNA synthesis, and transmission^96^. Thus, more research may be undertaken to determine if the BBTV-associated alphasatellite interacts with the host’s TFs to enhance viral pathogenicity in banana. The differential expression of these TFs in banana during BBTV may also affect the expression of genes involved in the biosynthesis of several hormones. As a possible consequence, the overexpression of genes involved in salicylic acid (*PR-1*) and jasmonic acid (*AOS*) signaling pathways and the downregulation of *IAA/AUX* for auxin perception and signaling, may be part of the BBTV’s attempt to subjugate the banana defense system. Moreover, our results also suggest that the invading BBTV pathogen also recruits the host SCF (SKP1-CUL1-F-box protein) ubiquitin ligase complex, which typically functions for JA signaling and defense responses, to enhance the degradation and ubiquitination of its pathogenesis machinery via ubiquitin/26S proteasome system^45^. Viruses achieve this mechanism via molecular mimicry in which the virus-encoded F-box protein interacts with the host SCF complex to attenuate plant antiviral defenses^45,97^.

Heat shock proteins, particularly HSP70, are essential components of plant immunity, taking part in both PTI and ETI responses^98^. However, viruses use the HSPs of the host for their transcription, translation, post-translational modification and cellular localization^99^. When compared to the resistant banana genotype, higher number of exclusively differentially expressed HSP genes was observed in the susceptible banana genotype (1 DEG in resistant, 8 DEGs in susceptible banana). Hence, rather than aiding in plant immunity^27^, the HSPs in banana are being exploited by BBTV for its own benefit. In response to viruses of resistant plants, the downregulation of HSP proteins was observed to restrict viral movement through plasmodesmata^100^. Therefore, silencing or inhibiting the expression of these HSP genes can contribute to preventing the progression of diseases caused by viral pathogens^101–103^.

### Differential transcriptomic analysis provided insights into resistance mechanisms of banana against BBTV

By comparing to the susceptible banana genotype, we were able to identify resistance genes that may contribute to the high degree of BBTV resistance reported in wild *M. balbisiana*^8^. Our work demonstrates that natural BBTV resistance found in the wild encompasses complex and multi-component mechanisms^104^. These include but are not limited to pathogen detection and response, action of phytohormones, reactive oxygen species (ROS), hypersensitive response (HR), and production of secondary metabolites.

Unlike the susceptible ‘Lakatan’ genotype, the resistant wild *M. balbisiana* was found to have a greater number of exclusively differentially expressed protein kinase genes, which may be involved in pathogen recognition and signaling, activating the overall resistance mechanisms in banana against BBTV. Intriguingly, after BBTV infection, gene expression of *WAK* was up-regulated in the resistant banana genotype but not in the susceptible banana genotype. It has been demonstrated that WAKs are capable of detecting damage-associated molecular patterns (DAMPs), which are made up of pectin and oligogalacturonide molecules released from the plant cell wall as a result of pathogen infection. These damage signals are subsequently sent by WAKs, which affects both plant development and defense^47,105,106^. Transgenic plants that overexpress the WAK gene have been shown to improve pathogen resistance^107,108^. Moreover, introgression of WAK gene in maize marker-assisted breeding programs enhances head smut resistance^109,110^. Hence, the WAK gene found in wild *M. balbisiana* warrants further functional and genetic investigations to establish its potential significance in the banana BBTV resistance breeding program.

BBTV-resistant wild *M. balbisiana* also uses phytohormone action to defend against the viral invasion. Salicylic acid (SA), jasmonic acid (JA), ethylene (ET), abscisic acid (ABA), nitric oxide (NO), cytokinins (CK), gibberellin (GA), auxin, and brassinosteroids (BR) are examples of hormones that act downstream of pathogen detection providing another layer of defense^104^. Among these hormones, genes involved in abscisic acid signaling and auxin transport were downregulated in the resistant genotype.

Restriction of further pathogen infection via HR, ROS and cell modification is apparently being implemented by the resistant banana genotype to protect itself against BBTV. HR is a frequent immune response of plants that causes programmed cell death in the areas of pathogen infection. This produces a containment zone to prevent pathogens from spreading, which is an effective method for biotrophs such as viruses^104^. The contribution of hypersensitive-reaction protein in plant basal resistance against *Rice stripe virus, Turnip mosaic virus, Potato virus X*, and the bacterial pathogens *Pseudomonas syringae* and *Xanthomonas oryzae* via *EDS1* and salicylic acid-dependent pathways was previously demonstrated in *N. benthamiana* and rice^111^. Moreover, ROS-induced oxidative bursts also cause programmed cell death and create an unfavorable host environment, limiting further pathogen survival and proliferation^112^. In addition to redox reaction-related DEGs, the genes involved in the biosynthesis of polyamines (putrescine and spermidine) were also differentially expressed in the resistant banana genotype. The role of polyamines in plant immunity is linked to the regulation of hydrogen peroxide (H_2_O_2_), a predominant ROS in plants, through the catabolism by amine oxidases^113^. Moreover, genes involved in cell wall organization were differentially expressed. Various researches on essential plant defense cell wall-associated proteins in differential susceptible and resistance responses in plants demonstrate that cell wall organization (loosening and/or tightening) can influence viral propagation^114^. As such, the differential expression patterns observed in the genes involved in the biosynthesis and modification of various cell wall components in the wild *M. balbisiana* make them key players in banana-BBTV interaction.

The resistant wild banana appears to be undergoing differential production of secondary metabolites in response to BBTV exposure, including terpenes, carotenes, and flavones as evidenced by the differential expression of genes for their biosynthesis. Plant secondary metabolites are involved in several biological processes, including innate immunity and defense signaling^115^. As an example, a terpenoid phytoalexin was reported to be involved in the basal response of *N. benthamina* against *Potato Virus X* catalyzed by terpenoid synthase 1 (TPS1)^116^, suggesting a potential role of the TPS DEGs in banana BBTV resistance. Additionally, cross-talk between flavones and salicylic acid reveals that flavone biosynthesis is involved in plant-pathogen interactions. Interestingly, wild banana downregulates the expression of *FNS*, which is homologous to Arabidopsis DOWNY MILDEW RESISTANT6 (*AtDMR6*) and establishes interaction among flavones, hormone action and pathogen attack^117^. Meanwhile, although the role of carotenoids in plant pathogen immunity has not yet been established, defense response of wild banana against BBTV may be attributed to the relationship of carotenoids to atypical ROS singlet oxygen (^1^O_2_) and jasmonic acid signaling pathway^118–120^.

Transcriptional reprogramming is promoted by TF activity at various levels of resistance, such as the expression of proteins related to basal resistance, the direct TF activity of activated receptor proteins, and the activation of TFs downstream of receptor initiation^104^. Upon BBTV exposure, the wild *M. balbisiana* induces or down-regulates the expression higher number of genotype-specific TFs (25 TFs) as compared to the response of the susceptible ‘Lakatan’ to the pathogen (13 TFs). The expression of all NAC TFs were all suppressed in defense of wild banana against BBTV. NAC TFs were overexpressed upon infection of susceptible rice genotypes to *rice stripe virus* (RSV) and *Rice tungro spherical virus* (RTSV) while a gene knockout of NAC provides rice with resistance against *Rice dwarf virus*. Hence, our result warrants further investigation into how the identified differentially expressed TFs specific to the resistant genotype be deployed in actual banana BBTV resistance breeding programs.

## CONCLUSION

For the first time, this current research sheds insights into the fundamental understanding of the host-dependent infection process of the most destructive banana viral disease as well as unravels the natural resistance mechanism found in wild bananas. Comparative transcriptomic analysis was able to identify candidate resistance mechanisms in the wild *M. balbisiana* that might have been lost during the process of cultivation of commercial banana varieties for outstanding fruit quality traits. Thus, the identification and incorporation of the resistance alleles from the wild banana genotypes into the existing banana varieties is critical to developing DNA markers for use in marker-assisted plant breeding programs. Moreover, the potential host factors that are essential to the successful invasion of BBTV in banana that this research provides can be targeted for genome editing (eg. CRISPR-Cas) towards development of new sources of resistance alleles. Hence, this paper laid the groundwork to further advance the status of molecular breeding programs essential to safeguard the banana industry from the threats brought about by BBTV.

## ACKNOWLEDGEMENT

This research was made possible through funds provided by the Philippine Department of Agriculture - Biotechnology Program Office (DA-BIOTECH) to the project entitled “DA-BIOTECH R1902: Fast-tracking the Development of BBTV-resistant Banana Cultivars through Modern Biotech Tools: Molecular Profiling towards Marker Development and Diagnostics (Phase I).” The authors gratefully acknowledge the Institute of Plant Breeding for the use of its facilities and equipment. The authors would also like to express their sincere appreciation for the technical and administrative assistance provided by Ronilo M. Bajaro, Wilermie Driz-Hernandez, Reina Esther S. Caro, and Rodelio R. Pia.

## Author Contributions

DVL, ANCM, RRG, GCL and FMDC contributed to the study conception and design. RNA extraction was performed by JDLN with supervision by ANCM. Preparation and treatment of experimental seedlings were conducted by JSM. LSG provided the germplasm materials. DVL performed the data analysis and interpretation, and wrote the original draft. All authors read and approved the final manuscript.

## Notes

### Competing Interest Statement

The authors have declared no competing interest.

